# Multiscale Spatial Mapping of Microbial Communities for Biotherapeutic Development

**DOI:** 10.1101/2025.03.31.646377

**Authors:** Prateek Sehgal, Andrea G. Shaw, Kevin Cutler, Bridget Letson, Gregory T. Booth, Gaetano J. Scuderi, Anjali Doshi, Peter Diebold, Hannah Bronson, Maya Trisolini, Unnati Dharaiya, Kevin Keomanee-Dizon, Seyi Jung, Jennifer Bagin, James Berleman, Aditya Bhalla, Kyle Jacoby, Steve Kujawa, Jackson Mintz, Pallavi Murugkar, Marguerite Prior, Keiramarie Robertson, Ashlie L. Burkart, Jason Pontin, Lee R. Swem, Iwijn De Vlaminck, Matthew P. Cheng, Philip Burnham, Hao Shi

## Abstract

Live biotherapeutic products (LBPs) are emerging as powerful tools to modulate the microbiome using well-defined microbial communities. Yet, designing, manufacturing, and delivering LBPs remains challenging, in part due to a lack of technologies capable of analyzing LBPs as complete, spatially organized consortia. Conventional sequencing-based methods lack sensitivity and specificity and do not provide critical spatial information. To address this, we present high-phylogenetic-resolution spatial mapping platform (HiPR-Map), a state-of-the-art spectral imaging technology that enables precise enumeration and spatial localization of microbial cells at species-level within complex communities. Through these advantages, HiPR-Map provides unique insights for LBP discovery and development. Applying HiPR-Map to an LBP designed to complement immune checkpoint therapy, we profiled over 1.8 million microbial cells engrafted in the murine gut. Our analysis revealed distinctive microbial spatial organization, underscoring the power of imaging-based microbiome profiling to optimize LBP design and characterization. This work highlights the transformative potential of spatial microbiome analysis for next-generation LBP development.

## INTRODUCTION

Microbes are intimately tied to maintaining human health, contributing to nutrient digestion, immune regulation, drug metabolism, and disease progression. Microbial therapies through the use of fecal material transplants (FMTs) have recently emerged as a promising approach to treat infectious diseases and various oncology indications^1–4^. However, FMTs face significant challenges related to reproducibility, safety, and manufacturing at scale, which limit their broader clinical adoption. As a result, there has been a shift towards the development of live biotherapeutic products (LBPs) as a standardized and scalable alternative to FMT-based therapies^5^. LBPs, which consist of a defined consortium of microbes, require precise characterization throughout their lifecycle—from manufacturing to clinical administration—to ensure product consistency, mechanistic understanding, efficacy, and safety.

Optimizing LBPs requires detailed insights into microbial composition, spatial organization, microbe-microbe interactions, and host-microbe interactions. Sequencing-based methods provide valuable compositional and functional insights in the analysis of LBPs. However, they are limited in capturing spatial organization, absolute abundances, and *in situ* microbial interactions. Specifically, sequencing approaches such as 16S amplicon sequencing and metagenomic sequencing require sample homogenization, which removes spatial information. Spatially-resolved sequencing methods retain this information but are limited in spatial resolution and sensitivity for microbes^6–8^. Furthermore, sequencing-based methods often struggle to identify rare taxa (<0.1%) and are prone to contamination in low biomass settings^9^. As a result, critical factors in LBP manufacturing such as purity and potency, in addition to key aspects of *in vivo* LBP analysis such as engraftment, strain interactions, and host-microbe dynamics, remain poorly understood, limiting the development of effective LBPs.

Recently, imaging-based methods have emerged as a powerful tool to overcome the limitations of sequencing^5,10–25^. These approaches enable microbiome analysis by revealing microbial spatial organization across varying length scales^11^. However, they are generally constrained by low phylogenetic resolution and multiplexity, restricting their ability to capture microbiome diversity. Despite recent advances in enhancing multiplexity^26^ and their vast potential in advancing LBP development, imaging-based profiling of microbial communities at the species level continues to be challenging.

Here, we present HiPR-Map for single-cell and spatially resolved microbiome profiling at the species level. HiPR-Map incorporates advances in probe design and image analysis that are marked steps forward from previous fluorescence *in situ* hybridization (FISH)-based assays – including the ability to resolve rRNA at single-nucleotide resolution and the ability to acquire data over large areas, spanning whole tissue sections. These advancements enable accurate analysis of microbiome composition, spatial organization, and cell state across the development cycle of LBPs.

## RESULTS

### HiPR-Map assay and comparison with sequencing

HiPR-Map uses spectral barcoding for highly multiplexed taxonomic labeling of microbes via the hybridization of probes to 16S ribosomal RNA (**Fig. 1a**). To demonstrate the sensitivity and specificity of HiPR-Map, we first designed a species-level probe panel targeting 15 diverse taxa representing varying degrees of taxonomic and phenotypic diversity, ranging from gram-positive, gram-negative, aerobic, and anaerobic bacteria, as well as a fungus and archaea (**Table S1**).

**Figure 1:**
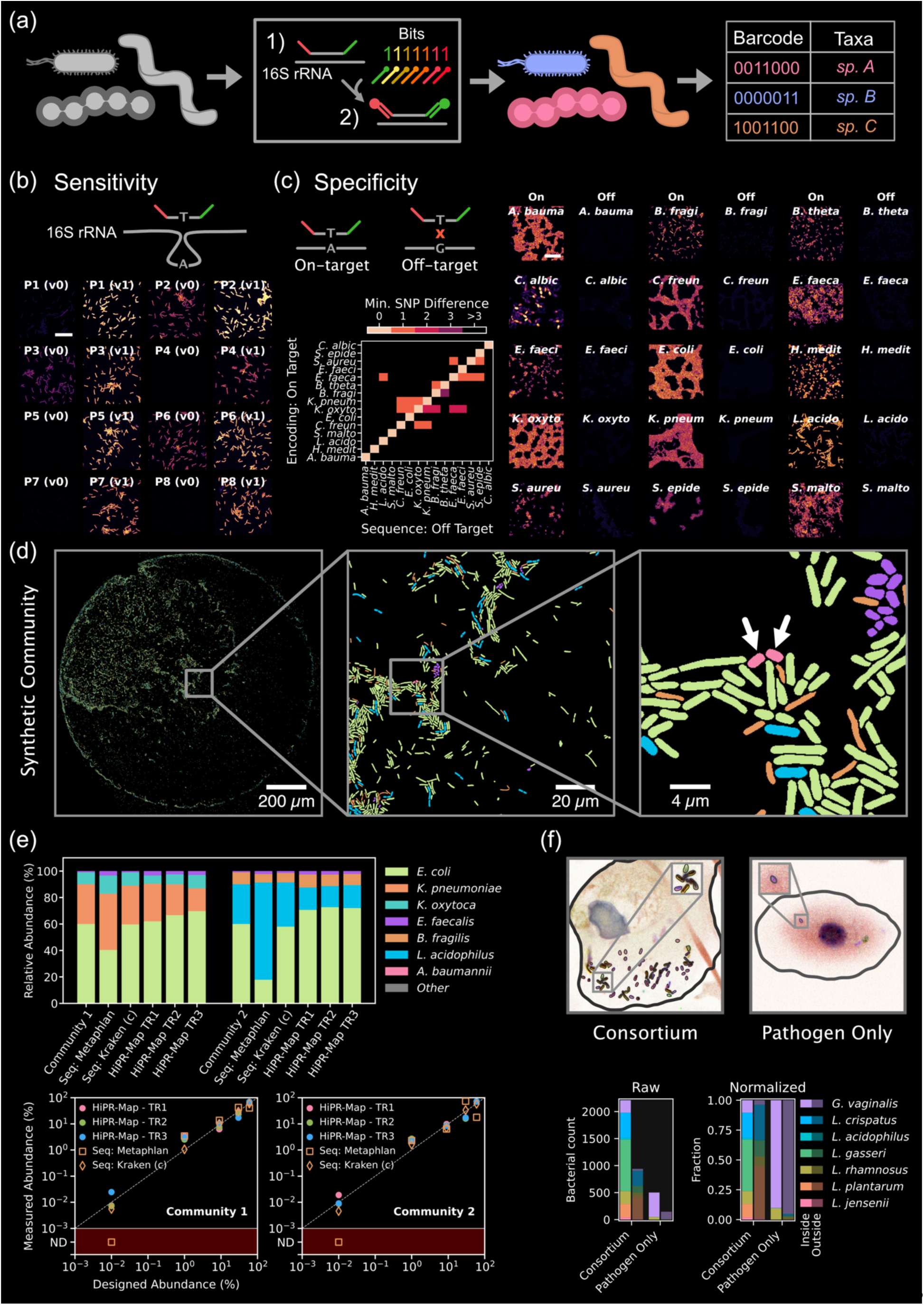
Technological advances of the HiPR-Map platform. (a) HiPR-Map platform utilizes a highly multiplexed labeling approach to fluorescently barcode microbial rRNA. Each encoding probe binds up to two fluorescent probes (readout probes), encoding up to two bits of information. Combinatorial addition of several encoding probes per unique encoding region on rRNA enables multi-bit barcoding of each taxon^26^. Hyperspectral data is collected via confocal imaging, and advanced image analysis is used to segment and classify microbiota. (b) HiPR-Map probe design achieves high sensitivity by enhancing ribosomal accessibility (schematic). HiPR-Map (labeled “v1”) shows significantly high sensitivity over HiPR-FISH^26^ (labeled “v0”), with all encoding probes generating bright and detectable signals. The intensity distribution is shown in **Fig. S2**. The scale bar represents 20 µm. (c) HiPR-Map probe design achieves high specificity by preventing encoding probes from binding to off-target sequences that differ by a single nucleotide from the on-target sequence (schematic). (Left) The heatmap shows the minimum single nucleotide polymorphism (SNP) difference for all encoding probes between members of a 15-species community. Several encoding probes only differ by a single nucleotide from the off-target sequences. (Right) Images demonstrate high specificity by comparing measured on-vs off-target signal for individual species. The fold-change is shown in **Fig. S3**. The scale bar represents 20 µm. (d) Full-scale imaging of a synthetic community generated from the 15-species pool and dried on a slide. Labeled cells, colored by their respective taxon identity, are shown in the images. The double inset shows HiPR-Map detecting *A. baumanii* cells present at a designed relative abundance of 0.01% in this community. (e) Two distinct communities were designed (see **Methods** for cell counting assay) and measured in triplicate using HiPR-Map. The relative abundance obtained from HiPR-Map was compared to metagenomic sequencing data processed using Kraken (with Bracken) and MetaPhlAn (see **Methods**), as well as to the designed abundance. (Top) Relative abundances on linear-scale, and (Bottom) Relative abundances on log-scale. “ND” refers to Not Detected. (f) (Top) Processed images show the adhesion *of G. vaginalis* and a Lactobacilli consortium to vaginal epithelial cells (VECs). (Bottom) The absolute and relative abundances of each taxon is shown for bacteria adherent to the VECs (“inside”) and those adherent to the plate surface (“outside”).

The HiPR-Map probe design strategy enhanced the accessibility of the target site on the ribosome, dramatically improving the sensitivity of all encoding probes, including those that were ineffective when designed using conventional approaches (**Fig. 1b**). This high sensitivity enabled precise discrimination between 16S sequences of different taxa, even when the target site differed by only a single nucleotide (**Fig. 1c**, heatmap). We validated the specificity of the assay by testing the full panel against each species and assessing off-target signal intensity. Across all 15 species, we observed negligible off-target signal relative to the on-target signal, with signal-to-noise ratios ranging from 10 to 100 (**Fig. 1c** & **S3**). This high sensitivity and specificity achieved with the probe design, staining, and imaging approach of HiPR-Map enabled accurate profiling of microbiome composition across diverse biological systems.

To evaluate the accuracy of HiPR-Map against sequencing-based microbiome profiling, we created two synthetic communities with distinct relative abundance (RA) profiles ranging from 60% to 0.01%. Both communities were assayed simultaneously using HiPR-Map and metagenomics (see **Methods** for details). **Fig. 1d** highlights the imaging scale achieved with HiPR-Map, demonstrating full-scale imaging of a ∼1.2 mm diameter droplet of a community on a microscope slide. In addition to precise cell counting, HiPR-Map enabled single-cell resolution mapping, allowing spatial localization of each microbial cell—a key advantage over sequencing-based approaches (**Fig. 1d**, zoom-in).

HiPR-Map achieved a detection limit of 0.01% RA, and critically, the RAs measured by HiPR-Map correlated well with the ground truth (designed) abundances for each synthetic community across all the replicates (**Fig. 1e**). In contrast, the abundances measured by sequencing depended strongly on the sequence alignment method used. For instance, MetaPhlAn consistently underestimated the abundance of *Escherichia coli* in both communities and overestimated the abundance of *Lactobacillus acidophilus* in Community 2. Furthermore, MetaPhlAn was unable to detect the lowest abundance taxa (0.01% RA) in either of the communities (**Fig. 1e**, bottom image, “ND” region). We hypothesized that this discrepancy was due to the broad database used for MetaPhlAn alignment. We therefore created a custom Kraken database comprised of only the taxa present in the designed communities, which more accurately predicted community composition and successfully detected low-abundance taxa. These findings underscore how sequencing-based measurements can vary depending on the alignment tool, particularly for low-abundance taxa.

To demonstrate the utility of HiPR-Map in low-biomass samples, we utilized a co-culture model commonly leveraged during LBP development to assess pathogen displacement on the skin^27^. Briefly, we co-cultured vaginal epithelial cells (VECs) with the pathogen *Gardnerella vaginalis* before introducing a synthetic *Lactobacillus* consortium containing six unique species (see **Methods**). After incubation, we quantified bacterial cells via HiPR-Map. We found that all species were adherent to the VECs (**Fig. 1f**) and the bacterial count for *Lactobacillus* species was significantly higher than *G. vaginalis.* Furthermore, the absolute count of *G. vaginalis* was suppressed in the presence of *Lactobacillus* consortium relative to the *G. vaginalis*-only controls, indicating successful adherence and partial displacement of the pathogen. HiPR-Map also detected microbes adherent only to the cell culture plate. Interestingly, normalizing these cell counts revealed distinct VEC-adherent (“inside”) and plate-adherent (“outside”) microbiota profiles. In contrast, sequencing would not be able to distinguish these populations, instead grouping all microbes together, distorting the apparent VEC-adherent microbiome composition. Collectively, these results demonstrate the utility of HiPR-Map to accurately profile microbiomes at unprecedented spatial and phylogenetic resolution.

### Spatial mapping of host-microbiome-food interactions to understand LBP engraftment

LBPs are being developed to enhance the efficacy of immune checkpoint inhibitor (ICI) therapy, but their success is hindered by challenges in characterization, manufacturing, and engraftment. To address this, we built an extensive library of microbial strains from ICI-responsive patients to develop LBPs that support treatments before, during, and after cancer therapy. We assembled a consortium of proprietary strains and evaluated their therapeutic potential in a germ-free mouse model, where mice were colonized with the consortium, followed by cancer implantation and subsequent ICI therapy. After treatment, mice were sacrificed, and their gastrointestinal (GI) tissues were harvested and fixed. We then used HiPR-Map to provide spatially resolved insights into microbial engraftment and distribution.

We profiled full tissue sections from the ileum, cecum, and colon (**Fig. 2a**). We imaged and classified over 1.9 × 10^5^, 1.4 × 10^6^, and 2.5 × 10^5^ microbial cells in the ileum, cecum, and colon, respectively. We observed extensive interactions of microbes with the host tissue, food particles, and other microbes (**Fig. 2b**). The species abundances were more similar between the cecum and colon (Spearman rank correlation, ρ = 0.95) than between the ileum and cecum (ρ = 0.80) or the ileum and colon (ρ = 0.78) (circles in **Fig. 2f**), showing that the ileum is compositionally different from the cecum and colon. For instance, species *BCA 057* is highly enriched in ileum (RA = 37.9%) but present at relatively lower abundances in cecum (RA = 2.5%) and colon (RA = 2.6 %). These findings support those previously reported^28,29^.

**Figure 2:**
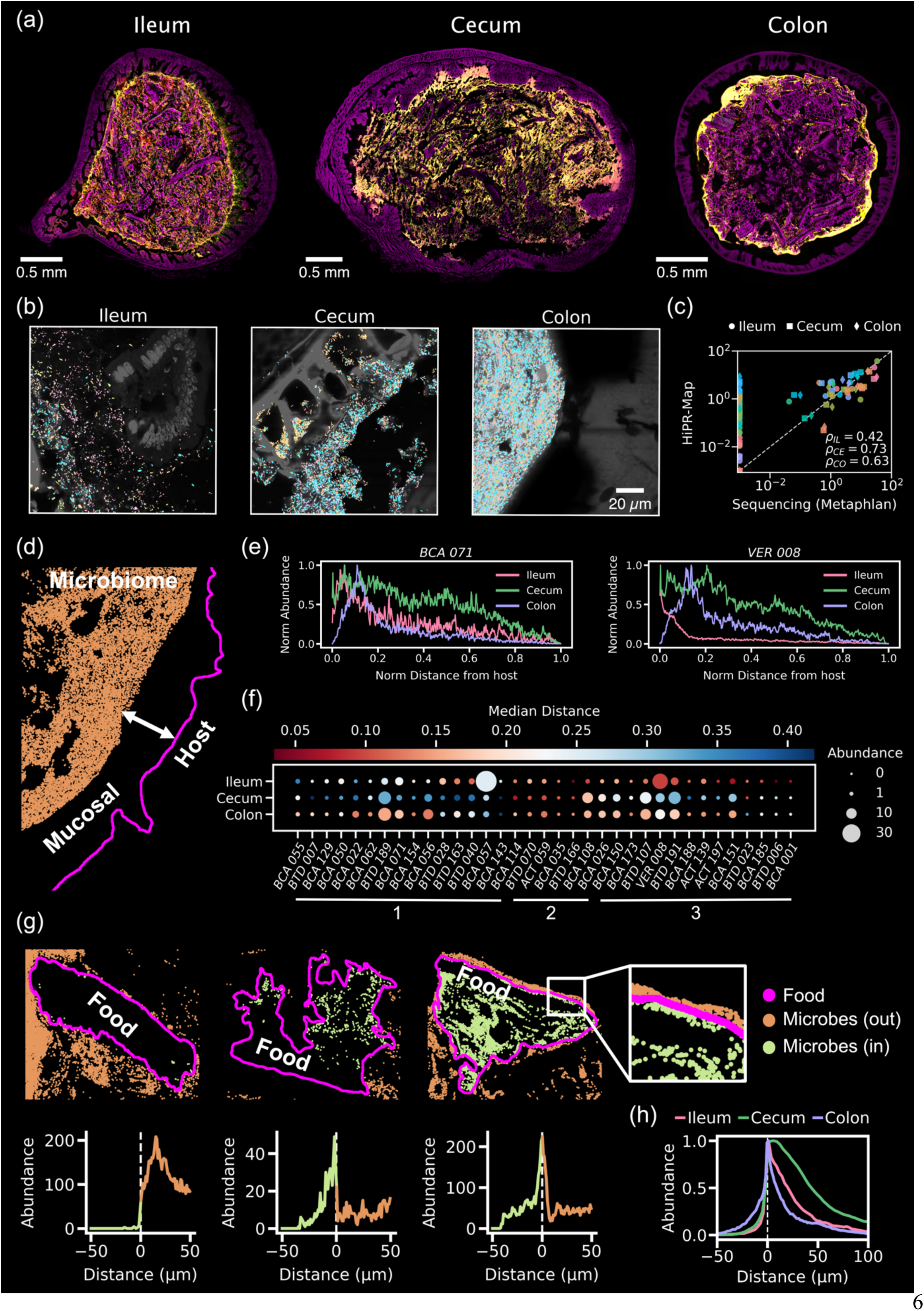
Whole tissue imaging and host-microbiome-food analysis. HiPR-Map’s strength lies in its ability to image across millimeter scales with sub-micron resolution. (a) Whole tissue hyperspectral imaging of the ileum, cecum, and colon of a single mouse that received consortium of proprietary strains designed to complement immune checkpoint therapy. Violet shows non-microbial material (food and host cells) and yellow-green-red shows the density of microbes. (b) Taxonomically identified microbes directly adjacent to the host epithelium (in ileum and colon), as well as within and surrounding the food particles (in cecum). The color legend for each taxon is shown in **Fig. S4**. (c) Comparison of relative abundances measured by HiPR-Map in tissue samples and sequencing of adjacent tissue sections. The color legend for each taxon is shown in **Fig. S4**. (Inset) The Spearman rank correlation between HiPR-Map and sequencing for the ileum (ρ_IL_), cecum (ρ_CE_), and colon (ρ_CO_). The correlations are calculated using taxa detected by both HiPR-Map and sequencing. The correlations using all taxa are shown in **Fig. S5**. (d) Centroids of individual microbes near host tissue, separated by mucosal barrier. These centroids were used to calculate the closest distance of each microbe to the host epithelium. (e) Histograms showing the abundance distribution of two taxa relative to the host boundary in the three tissues. Different taxa have different profiles depending on their environment. (f) Bubble plot depicting the median distance of each taxon from host tissue (normalized by maximum distance in that tissue) alongside their abundances in three tissue types. Bubble color represents median distance, while bubble size indicates relative abundance. The taxa on x-axis are grouped based on hierarchical clustering of median distances in all three tissue types. Three distinct groups (1, 2, and 3) are highlighted below x-axis. (g) (Top) Centroids of microbes inside (green) and outside (orange) the manually segmented food particles. The zoomed-in view highlights microbial distribution at the boundary of a food particle. (Bottom) Abundance profile of microbes within (negative) and outside (positive) the respective food particles. (h) Histogram showing the normalized abundance distribution of all microbes relative to the nearest food boundary in the ileum, cecum, and colon. Breakdown by each taxon is shown in **Fig. S6**.

We observed a strong correlation between species abundances measured by HiPR-Map and metagenomic sequencing of adjacent tissue across the ileum (ρ_IL_ = 0.42), cecum (ρ_CE_ = 0.73), and colon (ρ_CO_ = 0.63) (**Fig. 2c**). However, there were a few important exceptions. HiPR-Map detected several additional species that were not detected by sequencing (all-taxa rank correlations in **Fig. S5**), consistent with our observations from the synthetic communities (**Fig. 1e**). We attribute this to limitations of the sample preparation techniques used for sequencing, with inefficient cell lysis possibly leading to species dropout. Indeed, several of the missing species were gram-positive microbes which can be resistant to lysis^30^. Clearly, HiPR-Map accurately measures microbiome compositions relative to sequencing while simultaneously preserving the spatial organization of the tissue.

We next quantified the spatial distribution of microbes relative to the host by measuring the distance from each microbe to the nearest point on the host boundary (**Fig. 2d**). A few species in the LBP show preferential enrichment near the host boundary in different regions of GI tract, while other taxa show broader distribution into the lumen (**Fig. 2e**). To quantify these distribution patterns relative to the host, we calculated the median distance to the host boundary for each species in each of the tissues, hierarchically clustered these distances, and plotted them alongside their abundances (**Fig. 2f**).

We broadly found three groups with distinct spatial organization among the members of the LBP. Group 1 consisted of species that do not have preferential enrichment near the host tissue in all three regions of GI tract, Group 2 consisted of species that are enriched near the host in all three regions, and Group 3 consisted of species that are enriched near the host in ileum but not in cecum or colon (**Fig. 2f**). As an example, two taxa, *BCA 071* in Group 1 and *VER 008* in Group 3, highlight these characteristic differences between groups and across tissue. While both taxa are similarly distributed throughout the lumen in the cecum and colon, *VER 008* is enriched near host tissue in the ileum, while *BCA 071* is depleted. Furthermore, taxon *BCA 057,* which is highly abundant in the ileum, does not exhibit preferential enrichment near host cells. While there are several major differences in the studies, our findings support the claims made about spatial organization of the microbiome at the phylum level in a recent preprint^31^. For example, *Actinomycetota (ACT 059, ACT 139, ACT 197* in our study) prefers to be closer to the host cells, particularly in the small intestine (**Fig. 2f**). Since different cell types have different abundances along the GI axis, this information can be used to enrich LBPs with strains that engraft and grow closer to target host cells.

We also observed that a significant portion of all three tissue regions contained a high density of food particles. Using manual segmentation based on autofluorescence, we found that food particles occupied at least 26% of the ileum, 20% of the cecum, and 48% of the colon lumen area, with particles distributed throughout the lumen—suggesting an intimate relationship with the microbiome. The high density of food particles in the colon may partly reflect the timing of sample collection, as most food material had likely reached the colon by the time of mouse sacrifice. However, the relatively denser packing is also likely due to the reduced water content in the colonic lumen^32^. To further investigate the food-microbiome interactions, we profiled microbial organization both surrounding and within the food particles.

For each food particle, we measured the distances of all microbes from its boundary, assigning negative values to those within the particle, and generated comprehensive microbial distribution profiles. We observed food particles with various characteristics of microbial proximity, including those with microbes distributed within their boundaries and others with microbes surrounding their boundary (**Fig. 2g**). The total microbial distribution relative to the nearest food boundary suggests that microbes were preferentially enriched closer to the boundary of food particles in all three regions of the GI tract (**Fig. 2h**). This result is in contrast with the previous work in other model systems, which reported a general underrepresentation of microbes within 5–10 µm of food particles^5^. However, we did not identify any species that exhibited a distinct overall pattern in its proximity to food particles (**Fig. S6**). We also found that the fraction of microbes located within the food particles was higher in the colon (purple curve in **Fig. 2h**) than in the ileum or cecum, which is likely due to the colon’s higher food density and the increased homogenization of luminal contents down the GI tract^32^.

Interestingly, we observed several large clusters of species *BTD 040* nested within the pockets of some food particles (**Fig. 2b**, cecum). Those food particles were identified as wheat through visual comparison with a reference image of wheat bran (**Fig. S7**). This observation aligns with previous reports that members of the *Bacteroidota* phylum have a broad ability to ferment polysaccharides commonly found in wheat^33^. Further studies with controlled diets and microbial gavage could provide deeper insights into such food-microbe interactions. Overall, this data demonstrates the utility of HiPR-Map to unravel food-microbiome interactions with high spatial and phylogenetic precision.

### Spatial analysis of intra-species interactions

A further strength of HiPR-Map is its ability to quantify the spatial organization of microbes across multiple length scales, from single-cell resolution to mesoscale clustering. To demonstrate this, we first quantified microscale self-aggregation of cells belonging to the same species. This approach was inspired by the visual observation of micro-colonies formed by different species across tissue regions in our images (**Fig. 2b**). We analyzed the distribution of each species individually by applying the density-based spatial clustering method, DBSCAN^34^. We set a ∼2 µm radius as the neighborhood around each cell and required a minimum of four cells within this radius to define a cluster.

We observed that several species exhibited differential self-aggregation across regions of the GI tract (**Fig. 3a**). For example, *BTD 107* formed clusters in the cecum and colon but not in the ileum, *BCA 057* aggregated in the ileum but not in the cecum or colon, while *BTD 189* and *VER 008* formed clusters in all three regions. To quantify self-clustering, we calculated the clustering strength (C-strength) of each species, normalized by its absolute abundance, across all three regions (**Fig. 3b**, top). This analysis provided a direct representation of species-specific self-aggregation as observed in the images. We then assessed which self-clustering interactions were statistically significant. To do this, we generated 5,000 simulations, randomly assigning species identities to the existing cell distributions in each tissue type while preserving their proportional abundances. To determine significant self-clustering (*p* < 0.01), we calculated the clustering strength from these random simulations and used it to normalize the measured C-strength (**Fig. 3b**, bottom).

**Figure 3:**
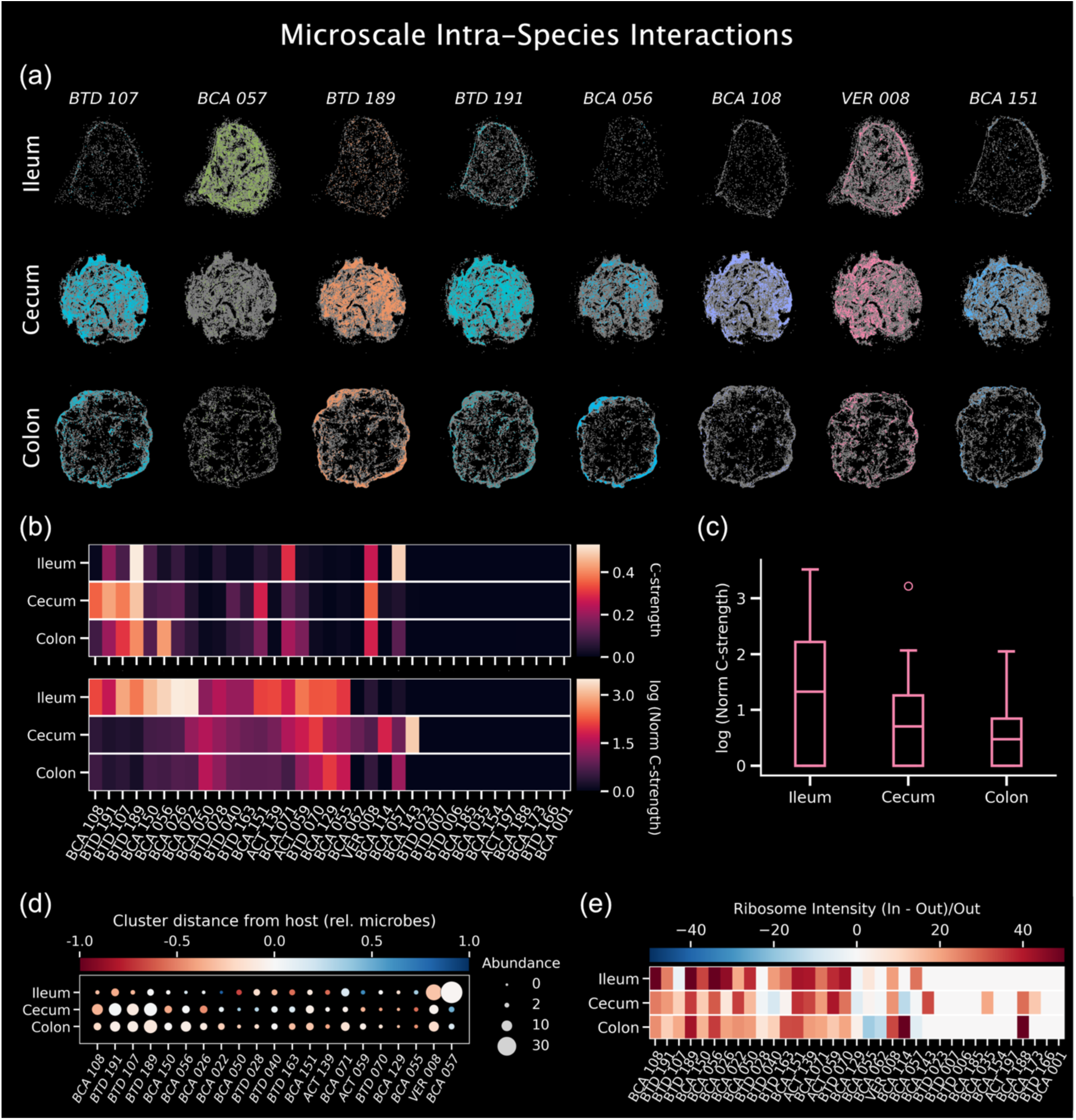
Measurement of intra-species interactions and ribosomal intensity. (a) Spatial distribution of self-clusters for several individual taxa throughout the GI tract. Colored regions indicate microbes that form clusters, while gray represents microbes that do not. Clustering is performed for each taxon using DBSCAN, with a ∼2 µm radius defining the neighborhood around each cell and a minimum of four cells required to form a cluster. (b) (Top) Self-clustering strength defined by cells that form cluster normalized by absolute abundance (C-Strength) for each taxon in each tissue. (Bottom) C-strength in top heatmap normalized by the C-strength measured from 5000 random simulations. Only significant values are shown (p < 0.01). Higher C-strength indicates a higher tendency to self-aggregate. (c) Normalized clustering strength aggregated across all strains in each region of the GI tract. (d) The median distance of identified clusters of a taxon from the host tissue, normalized by the median distance of all microbes of that taxon from the host tissue. The color of the disk represents distance and the size represents abundance. Only taxa that form clusters in all three regions of GI tract are shown for comparison. A value of −1 indicates that the taxon forms clusters closer to the host compared to its overall distribution, while a value of +1 indicates that the taxon forms clusters farther from the host relative to its overall distribution. (e) Comparison of ribosomal intensity between cells that are within clusters and those outside of clusters, analyzed by taxon and tissue type. Red indicates taxa where clustered cells have higher ribosomal intensity than non-clustered cells, while blue indicates the opposite. The color bar extrema are capped at –50% to +50% for visualization purposes. Taxa that do not form any self-clusters are assigned a zero value.

After normalization, we found that several species showed significantly enriched or depleted self-interactions. For example, *BCA 02*2 and *BCA 026* showed a higher tendency to cluster despite their lower abundances in ileum, while *BCA 057*, which was highly enriched in ileum, displayed fewer significant interactions over a random distribution. These results suggest that self-clustering is driven by two key factors: cell abundance and affinity between cells. Notably, when a lower-abundance taxon formed a self-cluster, it is more likely driven by intrinsic cell affinity rather than physical constraints of their abundance. Strikingly, we also observed that the normalized clustering tendency decreased along the GI tract (**Fig. 3c**), which is consistent with the increasing homogenization of fecal matter as it progresses through the gut^32^.

We measured the spatial positioning of self-clusters for each species relative to the overall distribution of all microbes from that taxon (**Fig. 3d**). We found that a few species, such as *BCA 050*, preferentially formed clusters closer to the host in the ileum, whereas *ACT 059* tended to cluster further away in the lumen. In the cecum and colon, cluster locations were largely driven by the overall distribution of each taxon, including for *BCA 050* and *ACT 059*. These results further underscore the differences between the ileum and the cecum or colon in host-microbiome interactions, aligning with the observations from previous studies^31,35^.

Lastly, we measured the ribosomal intensity of each species within self-clusters and compared it to that of non-clustered cells of the same species (**Fig. 3e**). Overall, we found that ribosomal intensity was higher in clustered cells than in those that did not form clusters. These results suggest that cells within clusters express more ribosomes, potentially indicating higher cell activity and division within these micro-colonies^16^. Consequently, our observations suggest that ribosomal intensity could serve as a biomarker for assessing cell health *in vivo*, aiding in the investigation of the mechanism of action of a given LBP.

### Spatial analysis of cross-species interactions

Next, we quantified spatial associations between different species at the microscale by calculating a region adjacency graph using the microbe segmentation mask (∼ 2µm length scale, **Methods**). We observed extensive cross-species association in all three tissue types (**Fig. 4a**, top row, **Fig. 4b**). We assessed significance by comparing the associations against randomized versions of the region adjacency graph and found that several associations were either significantly enriched or depleted (**Fig 4a**, bottom row). Most of the significant differential associations were tissue specific, but some associations were persistent throughout the intestine. *BCA 057*/*BCA 050* had a significant positive spatial association across the cecum and colon; *BCA 071*/*BCA 022* and *BCA 071*/*BTD* 070 had a significant positive association in the ileum and cecum. We did not observe persistent negative associations across different tissue types, suggesting that such interactions may be primarily influenced by the local gut environment. Nonetheless, identifying both positive and negative *in vivo* associations can inform novel interactions, which could in turn inform optimal co-culturing of species during LBP manufacturing.

**Figure 4:**
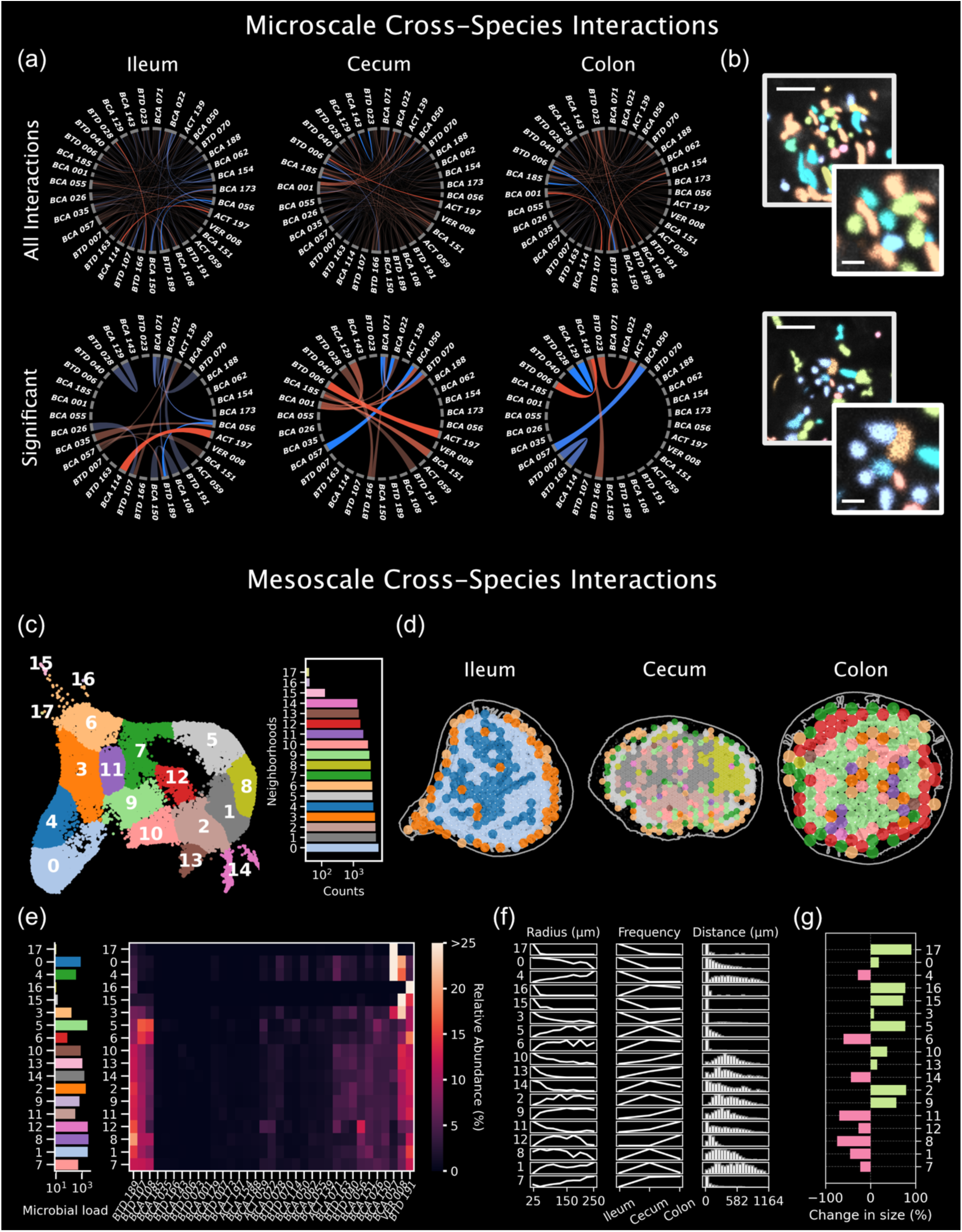
Cross-species interactions are frequent and show non-random structures. (a) Microscale cross-species interactions. Chord-style description shows absolute (top) and significant interactions (bottom; see **Methods**) across all measured taxa in each tissue. Here, self-interactions are excluded. Blue chords show positive associations, while red chords show negative associations. Brighter chords indicate stronger positive or negative interactions. (b) Taxonomically identified images show example inter-taxa interactions. The scale bars are 4 µm for the zoom-out and 1 µm for the zoom-in images. (c) Mesoscale cross-species interactions. For mesoscale interactions, disks ranging from radius 25 µm to 250 µm were generated and density of each taxon in each tissue was used to generate microbiome profiles. (Left) The disk profiles were projected into a 2D space using UMAP and a KNN and Leiden clustering approach was used to generate distinct neighborhoods. (Right) Frequency of each of these neighborhoods. (d) Distribution of neighborhoods projected onto radius-75 µm disks in each tissue. (e-f) Characteristics of the neighborhoods including microbial load (taxa per 10,000 µm^2^), relative abundances of different taxa, relative disk-size frequency, relative frequency per tissue, and distance from host. (g) Significance of neighborhoods over random distributions. Identity randomization was used to examine neighborhood reassignment (see **Methods**). Percentage change indicates the change in neighborhood frequency upon randomization. Strong positive values (green) indicate neighborhoods with strong, nonrandom spatial structures.

At the mesoscale (25–250 µm), we identified distinct neighborhood types based on their abundance profiles (**Fig. 4c**). These neighborhoods were defined using a grid of circular disks arranged in a hexagonal pattern, with spatial density calculated for each taxon in each disk (n = 52,559 disks). A UMAP projection of all neighborhood abundance profiles revealed 17 clusters, each representing a distinct neighborhood type (**Fig. 4c**, left). Notably, each tissue type contained a unique set of neighborhoods, highlighting the heterogeneity of spatial organization between these tissues at the mesoscale (**Fig. 4d**).

To further characterize these neighborhoods, we analyzed their compositional and spatial properties to identify large-scale spatial structures (**Fig. 4e-f**). For example, Neighborhood 12 (Nbd. 12) was enriched in the colon, and consisted of many taxa, with heightened relative abundances of *BTD 189* and *BCA 056*, and a reduced relative abundance of *BTD 191* compared with several other neighborhoods. This neighborhood was distributed away from host tissue, suggesting a distinctive ecological niche. In contrast, we identified low density neighborhoods with a high relative abundance of one particular taxon with strong tissue specificity, such as *BCA 057* (Nbd. 17) and *VER 008* (Nbd. 15) in the ileum, and *BTD 191* (Nbd. 16) in the cecum and colon. Importantly, these specialized neighborhoods were exclusively observed in direct proximity to host tissue. These findings were supported by an orthogonal self-aggregating cluster analysis, which identified similar spatial patterns (**Fig. 3b**).

To assess the significance of these spatial structures, we randomized microbial identities while preserving their assigned disk locations. The randomized profiles were projected into UMAP space, and clusters were reassigned accordingly. Comparing measured vs. randomized datasets, we found that many host-adjacent neighborhoods were significantly depleted, indicating strong spatial structuring. Conversely, some neighborhoods, such as Nbd. 8, increased in relative abundance, suggesting a more spatially random distribution (**Fig. 4g**). Our observations suggest that the significant mesoscale neighborhoods are indicative of the mechanism of action of LBPs and could serve as spatial biomarkers for predicting their therapeutic outcomes. Additionally, this analytical approach could be extended to identify predictive spatial biomarkers in feces. Overall, these *in vivo* cross-species interactions at various length scales offer invaluable insights into microbial cross-feeding, which is essential for optimizing co-culture manufacturing strategies for LBPs and evaluating their efficacy.

## DISCUSSION

The microbiome plays an important role in human health and is a promising target for the treatment of various diseases. While FMTs have demonstrated clear evidence of efficacy in infectious diseases and immuno-oncology^1–4^, there are limitations to their use. Realizing the clinical benefit of microbiome-based therapies requires reliable, reproducible, and scalable manufacturing of LBPs that recapitulate the diversity of FMTs. However, existing microbiome analysis tools, such as sequencing and culturing, have limited sensitivity and scalability for assessing LBPs and do not provide insights into spatial interactions. As a result, these methods restrict our ability to accurately characterize and understand LBPs throughout the discovery and development cycle.

In this work, we introduce HiPR-Map, an imaging tool for enumerating and spatially mapping microbiomes, and demonstrate its utility in LBP characterization, manufacturing control, and engraftment assessment. We show that HiPR-Map offers superior sensitivity compared to sequencing, which is critical for accurately assessing LBP engraftment in preclinical models. Additionally, HiPR-Map achieves the design of encoding probes that can distinguish sequences with single-nucleotide differences, which is necessary for microbiome mapping with species-level resolution. With these strengths, HiPR-Map enables detailed spatial analysis of complex microbial communities across different length scales, ranging from single cells to across the entire GI tract, revealing important insights that are otherwise lost in sequencing-based approaches.

Our analysis of therapeutic strain engraftment in the murine GI tract highlights the analytical power of the HiPR-Map assay. Importantly, we were able to recapitulate several findings reported by other groups. For instance, we observed that the ileum is both metagenomically and spatially distinct from the cecum and colon^31^, and that food particle density increases in the colon, consistent with water absorption by the epithelium^36^. Beyond validating previous observations, HiPR-Map uncovers novel insights into microbe-environment interactions, including relationships between microbial taxa, food particles, and host tissue. Notably, we identified several taxa enriched in regions adjacent to the mucosa. One such taxon, *VER 008*, a member of the *Verrucomicrobiota* phylum—known for its mucolytic capabilities^37^—was found to form aggregates in all tissue sections but was preferentially enriched near the host in the ileum. The cells in these aggregates exhibited higher rRNA content compared to non-aggregated cells, indicating higher activity and cell division in these aggregates. Additionally, *VER 008* does not show any statistically significant positive interactions with other taxa at the microscale. These lines of evidence suggest that *VER 008* preferentially grows in self-clusters without any significant interactions with other taxa. This vignette underscores the depth and resolution of HiPR-Map, which enables comprehensive, strain-level profiling of microbial communities within their native spatial context.

HiPR-Map insights can directly inform LBP product development, dosing strategies, and post-delivery monitoring of strain engraftment. By identifying which taxa colonize specific tissues, researchers can optimize strain selection based on therapeutic targets. For instance, a strain consistently found near immune structures like Peyer’s patches could be prioritized in a formulation to enhance immune system interactions. For more fine-grained resolution on host features, HiPR-Map’s spatial microbiome data can be combined with other spatial ‘omics measurements of the host cells to provide a comprehensive view of the host-microbiome interface. Integrating these tools into the development path of LBPs can accelerate LBP design iteration and provide mechanistic understanding of these therapies.

HiPR-Map’s high phylogenetic and spatial resolution also makes it a powerful tool beyond LBP development. It enables visualization of the microbiome in the tumor microenvironment, investigation of plant- and food-associated microbiota, and rapid culture-free microbial identification for infectious disease diagnostics. Taken together, HiPR-Map provides a transformative framework for precise microbial analysis and sets the stage for generating mechanistic insights into microbiome structure and function.

## METHODS

### Media preparation for cell culture

Bacterial and fungal media broths were prepared from dried broth mixes according to the manufacturer’s recommendations. Media for *H. mediterannei* was prepared using the ATCC Medium 1176: Halobacterium Medium protocol. Media used for anaerobic cultures was degassed in the anerobic chamber for a minimum of 48 hours prior to use. A full list of media used is listed in **Table S1**.

### Cell culture for synthetic communities

Cultured cells were inoculated from frozen ATCC stocks stored at −80°C into 2 mL of appropriate growth medium. Cultures were incubated overnight to reach the stationary phase. A 50 µL aliquot of the overnight culture was then transferred into 5 mL of fresh medium and incubated under optimal growth conditions. Each microbe was cultured in duplicate, and growth was monitored by measuring OD600 using a Biowave Cell Density Meter CO8000. Growth curves were plotted to assess microbial proliferation (**Fig. S1**). Cultures were harvested at mid-exponential phase for further analysis.

Microbial strain information, growth conditions, and optical density (OD) measurement at the time of harvesting are listed in **Table S1**. Once the cultures reached desired OD, two 2-mL aliquots of microbial suspension were transferred into separate tubes and centrifuged at 2,500 rpm for 10 minutes. The supernatant was discarded, and the pellets were processed differently. One pellet was resuspended in 100 µL of RNAlater for sequencing, while the second was resuspended in 750 µL of 4% paraformaldehyde (PFA) in 1X phosphate buffered saline (PBS) and incubated at room temperature for 15 minutes to allow fixation.

Following fixation, the cells were pelleted by centrifugation at 5,000 rpm for 5 minutes, resuspended in 1 mL of 1X PBS, and centrifuged again to remove residual fixative. This centrifugation and resuspension step was repeated once more to ensure complete removal of the fixative. The final pellet was then resuspended in 100 µL of 50% ethanol (EtOH) and stored at −20°C until further use.

*Bacteriodes spp*. were processed similarly, but due to the larger pellet size, cells were stored in 200 µL of RNAlater for one aliquot and 200 µL of 50% EtOH for the other.

### Probe panel construction

In this study, we leveraged our proprietary probe design pipeline to develop three distinct probe panels: (1) a 15-taxa synthetic community panel, (2) a 7-taxa *Lactobacillus*-*Gardnerella* consortium panel, and (3) a 35-taxa biotherapeutic consortium panel. All three panels were developed to barcode taxa at the level of species as defined by NCBI. Following design, probes were ordered from Integrated DNA Technologies (IDT) at 200 µM in IDTE pH 8.0 buffer. Each probe encoding a unique region was mixed at equal concentrations per unique encoding region as described previously^26^.

Following pooling, probes were concentrated via ethanol precipitation. 1X volume of the probe pool was combined with 0.1X volume of Sodium Acetate (pH 3.0) and 3.3X volume of 100% ethanol. The mixture was stored at −20°C overnight, before centrifugation at max speed (14,000 RPM) at 4°C. The supernatant was discarded, and the pellets were washed once with ice-cold 70% ethanol. The pellets were dried and resuspended in Tris-EDTA (TE) buffer.

### Synthetic community assay

Two microbial communities with varying abundances were designed by mixing a selection of 15 species. Communities assembled from fixed stocks were analyzed using HiPR-FISH, while those derived from RNAlater stocks were processed through metagenomic sequencing, as detailed below. To ensure accurate mixing ratios, cell counts from both fixed stocks and RNAlater stocks were measured using our proprietary cell counting assay, described below.

The HiPR-Map assay for synthetic communities was performed by drying three droplets of each community onto a glass slide. The samples were then lysed using lysozyme, followed by overnight encoding hybridization, a wash with wash buffer, readout hybridization, and a final wash. The slides were mounted in ProLong™ Glass Antifade Mountant for imaging. Details of the buffers used are provided in a previous study^26^. The communities were spectrally imaged on a confocal microscope as described below. The hyperspectral data was processed through our proprietary image analysis pipeline to decode the taxonomic identity of individual microbes.

### Cell counting assay

Fixed microbial cultures were suspended in 80% glycerol at the desired density for counting. RNAlater stocks were processed similarly, with the anaerobic microbes prepared in an anaerobic chamber. Following suspension, each sample was loaded into a Countess™ Cell Counting Chamber Slide, with equal volumes placed in both sides of the chamber. The chamber was sealed with epoxy (3M Scotch-Weld Epoxy Adhesive DP110 Translucent) and allowed to dry for approximately 15 minutes before centrifugation at 700 × g for 30 minutes. Images were acquired using a phase contrast Motic AE2000 microscope equipped with a digital camera (Moticam A16). Each sample was imaged at 40X magnification, systematically sampling different regions of the counting chamber.

### Metagenomics sequencing for synthetic community

Genomic DNA (gDNA) was extracted from synthetic communities stored in RNAlater (ThermoFisher, AM7024) using the ZymoBIOMICS DNA Microprep Kit (Zymo, D4301). Metagenomic sequencing libraries were prepared from synthetic community gDNA using Nextera XT DNA Library Preparation Kit (Illumina, FC-131-1096) and sequenced on Illumina’s MiniSeq system using a 300-cycle high output kit, generating approximately 8 million paired reads per community.

Raw fastq files were processed as follows. Reads aligning to the host genome were identified and removed using hostile^38^ with default parameters (human genome: human-t2t-hla), mouse genome: GCF_000001635.27_GRCm39). Non-host reads were then trimmed to remove adapter sequences and deduplicated using fastp^39^ with default parameters with the “--dedup” argument. With the remaining reads, the relative abundances in each community were determined using MetaPhlAn4^40^ with default parameters with their marker gene database (mpa_vOct22_CHOCOPhlAnSGB_202212). For comparison, relative abundances were also determined using Kraken2^41^ and Bracken with default parameters and a custom database built containing only genomes for the taxa that are part of the communities.

### Image acquisition

All imaging was performed on a Zeiss LSM 880, a high-performance inverted laser scanning confocal microscope. Spectral data were acquired in Lambda mode using 405, 488, 514, 561, 594, and 633 nm laser lines. A 63X oil objective was used to generate 135 µm × 135 µm fields of view. A line-averaging collection strategy was applied to reduce noise, and digital images were captured at 16-bit resolution. Zeiss’s Definite Focus module was used to maintain a consistent focal plane across the imaging surface. Custom scripts and Zen Black Macro Module were used to implement automated tilescan acquisition. All images—across fields of view and excitation channels—were stored as .czi files for downstream analysis.

### Segmentation and spectral classification

Segmentation was performed using multiple custom Omnipose^43^ models. A model for cell counting under phase microscopy was developed to achieve high accuracy with our optics and as compared to previous bact_phase_omni model. Three models were trained for segmenting bacteria imaged using confocal fluorescence under three sample preparation scenarios: (1) *in vitro* bacterial cultures deposited on slides (2) native bacteria within intestinal tissue sections, and (3) bacteria co-cultured with mammalian epithelial cells. Each of these models attained much higher precision and background suppression as compared to the previous best-in-class model, bact_fluor_omni. A final model was trained to segment host epithelial cells using background fluorescence, obtaining far better segmentation than available generalist models (*e.g.*, cyto2_omni). Training data for each model was curated manually using Napari.

Following segmentation of microbes, deconvolution of spectral data and classification of microbes were performed using our proprietary classification algorithm. Briefly, the spectrum of each segmented object was measured and matched to the closest spectrum in the training dataset, which represents all possible barcodes within a given consortium. This mapping process determined the taxonomic identity of each microbe.

### Co-culture assay with vaginal epithelial cells

Human vaginal epithelial cells (VECs) (Lifeline Cell Technologies catalog #FC1130) were cultured in 16-well chambers coated with Geltrex using ReproLife media (Lifeline Cell Technologies #LL-0068) until 70% confluency. *Lactobacillus* species (*L. crispatus, L. jensenii, L. gasseri, L. plantarum, L. rhamnosus, L. acidophilus*) and *Gardnerella vaginalis* were cultured in their respective media, as highlighted in **Table S1**, and grown to an optical density of 0.6 before co-culturing with VECs. Experimental conditions included *G. vaginalis*-only controls and a sequential inoculation of *G. vaginalis* followed by a mixed Lactobacilli consortium. Each condition was performed in four replicates. 25 million bacterial cells resuspended in antibiotic-free VEC media were utilized to inoculate the VEC for each condition. Slides were centrifuged at 150 × g for 5 minutes to promote bacterial adherence and then incubated for 1 hour at 37°C with 5% CO2. Wells were washed three times with antibiotic-free VEC media and fresh media either with or without mixed Lactobacilli consortium was added. Slides were again centrifuged at 150 × g for 5 minutes and incubated for 1 hour at 37°C with 5% CO2. Slides were washed again, fixed with 4% PFA in Dulbecco’s phosphate buffered saline (DPBS) for 15 min at room temperature, washed twice with DPBS, and stored in 70% ethanol at 4°C. Slides were then processed through HiPR-Map assay.

### Therapeutic consortium and mouse study

Isolated strains were streaked on specific agar and confirmed using colony morphology and MALDI-TOF spectrometry. These colonies were then cultured in anaerobic broth at 37°C until they reached necessary growth levels. Both the total cell count and viability were assessed before transferring the precultures into main cultures, which were similarly incubated. After achieving sufficient growth, a sterile glycerol solution was added for a final concentration of 25% v/v, and about 0.9 mL was aliquoted into cryovials. Once removed from anaerobic conditions, the vials were immediately stored at ≤ −65°C to complete the research cell bank (RCB) banking process.

To generate candidate therapeutic consortia for oral gavage, RCBs were mixed to achieve roughly equal cell counts for each taxon. First, an equal number of RCB vials of each strain comprising the candidate consortium were anaerobically pooled into a sterile container. The average viable cell count (VCC) of the pooled material was approximately 6 × 10^8^ cells/mL. After gentle resuspension, the consortia mixtures were aliquoted into 0.9 mL aliquots for cryopreservation at ≤ −65°C. On the day of administration (see below), the input candidate consortia were thawed and immediately introduced by gavage as a cell suspension.

The mouse experiment was performed as part of a larger study at the Centre Hospitalier de l’Université de Montréal (CHUM). Briefly, germ-free mice were inoculated with the consortium via oral gavage 15 and 14 days prior to the subcutaneous injection of MCA205 cancer cells. At days 6, 9, 12, and 15 following tumor cell injection, mice were injected with anti-PD-1 antibody intraperitoneally, tumors were measured, and feces were collected. Seventeen days after MCA205 injection, the mice were euthanized for rapid organ harvesting and tissue processing.

The GI tract was harvested from beneath the stomach to the rectum and dissected into the small intestine, cecum, and colon. Each segment was promptly rinsed with ice-cold 1X PBS to clear any residual blood. Next, the tissues were submerged in freshly prepared 10-15 mL of 4% PFA in a flat-bottom vessel and fixed for 16-24 hours at 4°C. The tissues underwent a sucrose gradient treatment, first in 15% sucrose in 1X PBS for 6-12 hours, followed by 30% sucrose until each tissue piece sank to the bottom of the container, indicating adequate penetration. Following sucrose treatment, the tissues were rinsed once more in ice-cold 1X PBS and embedded in OCT with liquid nitrogen. Frozen tissues were transferred on dry ice to −80°C storage. For analysis of the microbiome in this paper, we chose a mouse with the lowest tumor burden with the drug treatment cohort.

### Metagenomics sequencing for mouse study

To generate a metagenomics profile for each tissue, OCT-embedded sectioning was performed on a cryotome (Epredia Cryostar NX50). For each tissue, five 20 µm sections were transferred to 1.5 mL Eppendorf tubes for DNA extraction. Genomic DNA was then extracted from tissue sections using the Quick-DNA FFPE Miniprep kit (Zymo, D3067). Library preparation (Illumina DNA Prep kit, also known as Nextera Flex) and sequencing (Illumina, paired-end) were performed by Zymo Research – Next Generation Sequencing Service. We requested 20 M, paired-reads at 2×150 bp per sample. Reads were processed as described above using MetaPhlAn.

### Tissue sectioning and processing

The fixed frozen tissues were embedded in OCT at CHUM. Tissue sections of 10 µm thickness were obtained using a cryotome (Epredia CryoStar NX50) and mounted on SlideMate^TM^ Plus slides (Epredia). The slides were then incubated at 60°C for 5 minutes, followed by re-fixation with 4% PFA for 10 minutes. After re-fixation, the sections were washed with PBS and permeabilized in 70% ethanol at −20°C for 3 hours. The slides were then processed using the HiPR-Map assay.

### Host segmentation for tissue images

The host-lumen boundary in tissue section tilescans was annotated using a combination of automated and manual segmentation. Automated segmentation was performed using a pre-trained U-Net from Segmentation Models PyTorch, fine-tuned on hand-annotated data. Post-processing steps included binary hole-filling to address small gaps and the removal of very small stray objects. Final refinements were made through manual annotation in Napari.

### Host-microbiome-food analysis

Food particles in tissue sections were annotated manually in Napari using their autofluorescence, prioritizing larger particles with distinct boundaries and morphologies distinguishable from microbial clumps or other debris. Automated post-processing (binary hole-filling and removal of stray pixels) was applied to the annotation masks to ensure clean boundaries.

To validate food type, we ground commercially available wheat bran (One in a Mill brand) with a mortar and pestle for 2-3 minutes. Particles were filtered between a commercial size 30 mesh and size 60 mesh, such that very large and very small particles were filtered out. A portion of the remaining particles were then embedded in Neg50 medium in a CryoTek and frozen on dry ice followed by storage at −80°C. 10 µm sections were cut and placed onto microscope slides, then mounted with a thin layer of ProLong™ Glass Antifade Mountant. Spectral images were acquired on a Zeiss LSM 990 with a 20X and 63X objectives. Spectral images were combined with a custom-built function to generate the RGB projection shown in **Fig. S7**.

### Spatial association analysis comparison with identity randomization

To enable spatial association analysis, a region adjacency graph (RAG) was calculated for all classified cells in each FOV by expanding the segmentation masks. The RAG is a weighted undirected graph where each node is a classified cell, and each edge is a contact between any two classified cells. The weight of the edge is the centroid-to-centroid distance between the two nodes connected by the edge. The spatial association network is calculated by filtering out any edges in the RAG that has a weight of more than 35 pixels (2.45 µm). To calculate a taxon-level spatial association matrix measured from each FOV (mSAM), each node in the spatial association network is labeled by its assigned species, and each edge is assigned an edge type defined by the species identity of the two nodes connected by the edge. The SAM is then calculated as a 2D histogram of the edge types present in each FOV. To calculate a randomized SAM (rSAM), the species identity assigned to each node was randomly shuffled. For each FOV, the rSAM is calculated as the average of 1000 random simulations. This approach preserves the spatial and species abundance constraints as measured in the physical sample and only randomizes the spatial association between different cells. The (rSAM) is calculated the same way as mSAM described previously. To visualize the spatial association matrices, we calculated a fold change for each taxon-taxon pair using *FC_ij_* = log_2_[(*mSAM_ij_* + 1)/(*rSAM_ij_* + 1)]. To assess statistical significance of the spatial associations, the mSAM values across all fields of view measured from a given tissue (ileum, cecum, or colon) were compared to the rSAM values generated from the same tissue using an independent t-test. The significance threshold was corrected for multiple hypothesis testing using the Bonferroni criteria. The number of hypotheses being tested in our case is *n*(*n* − 1)/2, where *n* is the number of species in the LBP consortia.

### Neighborhood composition analysis generation

Neighborhoods were constructed by randomly seeding circles of predefined radii (25, 50, 75, 100, 125, 150, 175, 200, 225, and 250 µm) and iteratively packing adjacent circles to completely cover the tissue. Each circle was assigned a unique identification number, and a profile of all centroids corresponding to classified microbial objects within each circle and internal to the host boundary was generated. For each circle, the taxonomic identities of the microbes were recorded, and a count vector representing the abundance of each taxon was generated. To normalize for area, the count vector was divided by the circle’s area (µm²), yielding a taxon-density metric. Circles with a summed taxon density below 1 cell per 1000 µm² were excluded from the dataset. This process was repeated for five independent trials per circle size across three tissue samples, resulting in a total of 52,559 circles.

To generate clusters and define microbiome neighborhoods (i.e., groups of similar circles), a Uniform Manifold Approximation and Projection (UMAP) approach was employed using the umap-learn Python package (version 0.5.7) with the following parameters: n_neighbors = 100, random_state = 42, n_components = 2, min_dist = 0.05, and metric = ‘cosine’. Following dimensionality reduction, a k-nearest neighbors (KNN) graph (k = 1500) was constructed, and Leiden clustering was applied to partition circles into distinct groups. This clustering process yielded 17 microbiome neighborhoods, each characterized by its microbial load (classified microbes per unit area) and taxon-specific density profiles.

To evaluate the systematic nature of the spatial structure, randomization of taxonomic assignments within circles was performed. Specifically, taxonomic identities were reassigned across microbial centroids within each tissue while preserving global taxon abundances and centroid positions. Each randomization was conducted 10 times per circle to mitigate bias. The UMAP transform described above was then applied to project the randomized circles into the original UMAP space. A KNN-based lookup table was used to assign each randomized circle to a corresponding microbiome neighborhood, allowing for comparison between observed and randomized spatial structures.

## Declarations

Kanvas Biosciences, Inc. has filed patent applications for several of the technologies described in this work. AB and JP are members of the board of directors for Kanvas Biosciences.

