## Supplementary Information for "Multiscale Spatial Mapping of Microbial Communities for Biotherapeutic Development"

### Table of Contents:


| | |
|---|---|
| **Table S1:** Cell culture information for synthetic community experiments and cell adherence assays. | **Pg 2** |
| **Figure S1:** Plots of individual growth curves for all synthetic community strains. | **Pg 3** |
| **Figure S2:** Distribution of intensities from Fig 1b. | **Pg 4** |
| **Figure S3:** Quantification of HiPR-Map specificity from Fig 1c. | **Pg 4** |
| **Figure S4:** Color legend of strains used in the consortium for host-microbiome-food interactions. | **Pg 5** |
| **Figure S5:** Abundances measured by HiPR-Map and sequencing in tissue samples. | **Pg 5** |
| **Figure S6:** Distribution of microbes in and around food particles. | **Pg 6** |
| **Figure S7:** Reference images of wheat bran. | **Pg 6** |

**Table S1: Cell culture information for synthetic community experiments and cell adherence assays.** Growth conditions (i.e. atmosphere, medium, and temperature) are listed for each microbe. The OD600 at time of harvest indicates when samples were fixed during their exponential phase.

| Experiments | Species | ATCC | Gram | Atmosphere | Medium | Temp (°C) | OD at time of Harvest |
|---|---|---|---|---|---|---|---|
| Synthetic community | *Acinetobacter baumannii* | 19606 | Negative | Aerobic | Bacto™Tryptic Soy Broth Soybean-Casein Digest Medium (BD Difco #236950) | 37 | 0.79 |
| Synthetic community | *Bacteroides fragilis* | 25285 | Negative | Anaerobic | Difco™ Reinforced Clostridial Medium (BD Difco #218081) | 37 | 0.65 |
| Synthetic community | *Bacteroides thetaiotaomicron* | 29148 | Negative | Anaerobic | Difco™ Reinforced Clostridial Medium (BD Difco #218081) | 37 | 0.67 |
| Synthetic community | *Candida albicans* | 14053 | NA (Fungus) | Aerobic | Difco™ YM Broth (BD Difco #271129) | 25 | 0.97 |
| Synthetic community | *Citrobacter freundii* | 8090 | Negative | Aerobic | Difco™ Nutrient Broth (BD Difco #234000) | 37 | 0.9 |
| Synthetic community | *Enterococcus faecalis* | 29212 | Positive | Aerobic | Bacto™ Brain Heart Infusion (BD Difco #237500) | 37 | 0.62 |
| Synthetic community | *Enterococcus faecium* | 19434 | Positive | Aerobic | Bacto™ Brain Heart Infusion (BD Difco #237500) | 37 | 0.87 |
| Synthetic community | *Escherichia coli* | 25922 | Negative | Aerobic | Bacto™Tryptic Soy Broth Soybean-Casein Digest Medium (BD Difco #236950) | 37 | 0.8 |
| Synthetic community | *Haloferax mediterranei* | 33500 | NA (Archaea) | Aerobic | ATCC Medium 1176: Halobacterium Medium | 42 | 0.82 |
| Synthetic community | *Klebsiella oxytoca* | 49131 | Negative | Aerobic | Bacto™Tryptic Soy Broth Soybean-Casein Digest Medium (BD Difco #236950) | 37 | 0.73 |
| Synthetic community | *Klebsiella pneumoniae* | 13883 | Negative | Aerobic | Difco™ Nutrient Broth (BD Difco #234000) | 37 | 0.87 |
| Synthetic community | *Lactobacillus acidophilus* | 4356 | Positive | Anaerobic | BD Difco™ Lactobacilli MRS broth (BD Difco #288130) | 37 | 0.64 |
| Synthetic community | *Staphylococcus aureus* | 25923 | Positive | Aerobic | Bacto™Tryptic Soy Broth Soybean-Casein Digest Medium (BD Difco #236950) | 37 | 0.6 |
| Synthetic community | *Staphylococcus epidermidis* | 12228 | Positive | Aerobic | Difco™ Nutrient Broth (BD Difco #234000) | 37 | 0.68 |
| Synthetic community | *Stenotrophomonas maltophilia* | 13637 | Negative | Aerobic | Difco™ Nutrient Broth (BD Difco #234000) | 30 | 0.66 |
| Cell adherence | *Gardnerella vaginalis* | 14018 | Variable | Anaerobic | ATCC Medium 1685: NYC III medium | 37 | 0.6 |
| Cell adherence | *Lactobacillus acidophilus* | 4356 | Positive | Anaerobic | BD Difco™ Lactobacilli MRS broth (BD Difco #288130) | 37 | 0.6 |
| Cell adherence | *Lactobacillus crispatus* | 33820 | Positive | Anaerobic | BD Difco™ Lactobacilli MRS broth (BD Difco #288130) | 37 | 0.6 |
| Cell adherence | *Lactobacillus gasseri* | 33323 | Positive | Anaerobic | BD Difco™ Lactobacilli MRS broth (BD Difco #288130) | 37 | 0.6 |
| Cell adherence | *Lactobacillus jensenii* | 25258 | Positive | Anaerobic | BD Difco™ Lactobacilli MRS broth (BD Difco #288130) | 37 | 0.6 |
| Cell adherence | *Lactobacillus plantarum* | 14917 | Positive | Anaerobic | BD Difco™ Lactobacilli MRS broth (BD Difco #288130) | 37 | 0.6 |
| Cell adherence | *Lacticaseibacillus rhamnosus* | 7469 | Positive | Anaerobic | BD Difco™ Lactobacilli MRS broth (BD Difco #288130) | 37 | 0.6 |

**Figure S1: Plots of individual growth curves for all synthetic community strains.** Blue line represents the reference culture to measure log-phase while yellow line represents the culture that is used in the HiPR-Map experiments. The larger red dot indicates the latest timepoint of harvesting.

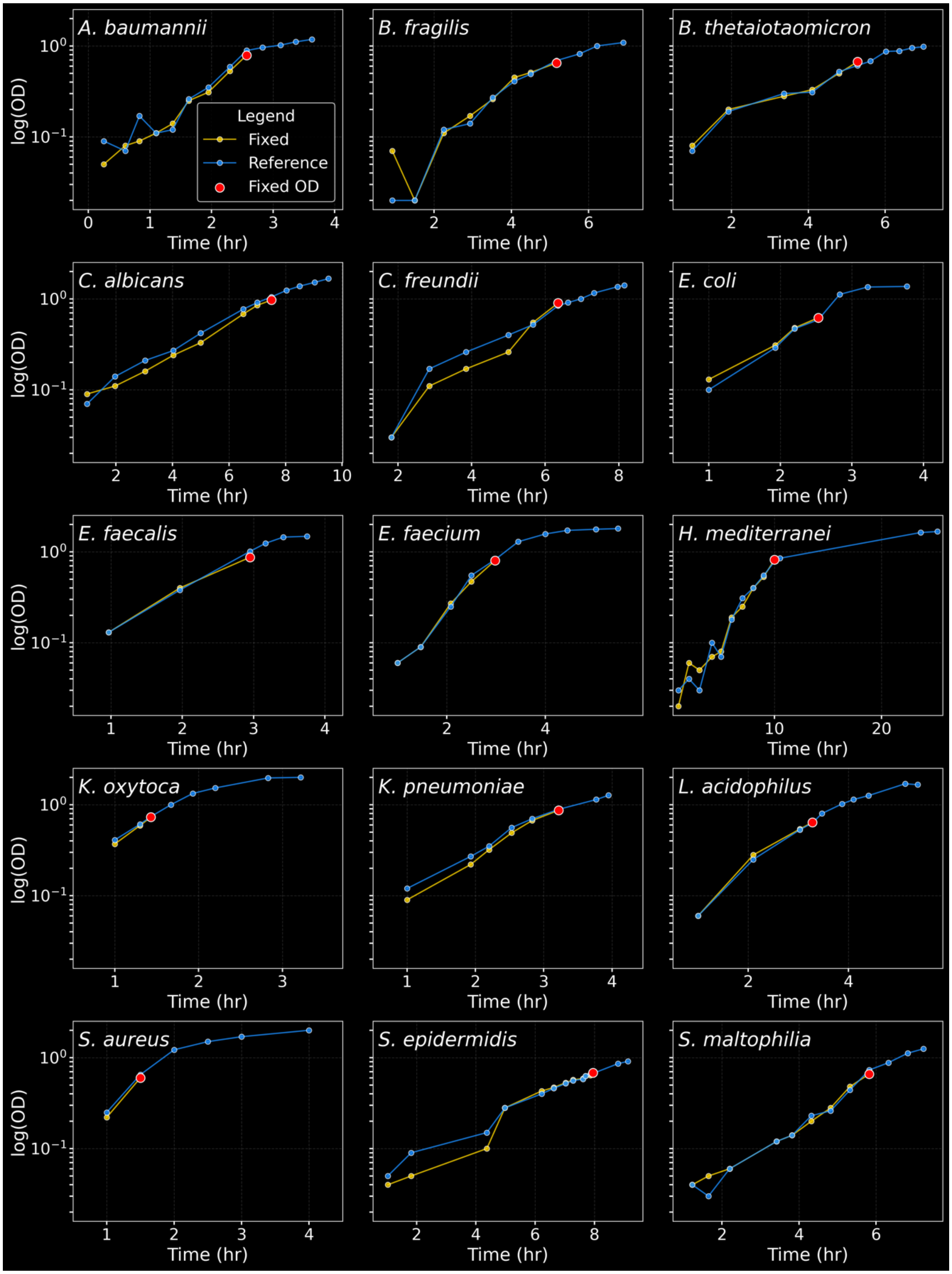

**Figure S2: Distribution of intensities from Fig 1b.**

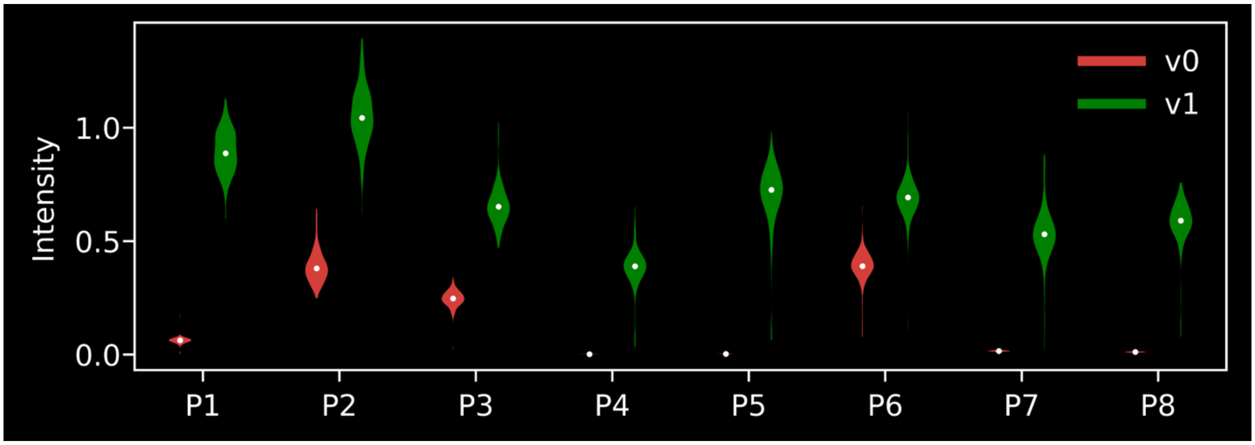


**Figure S3: Quantification of HiPR-Map specificity from Fig 1c**. Bars represent the fold change between on-target and off-target signal intensities.

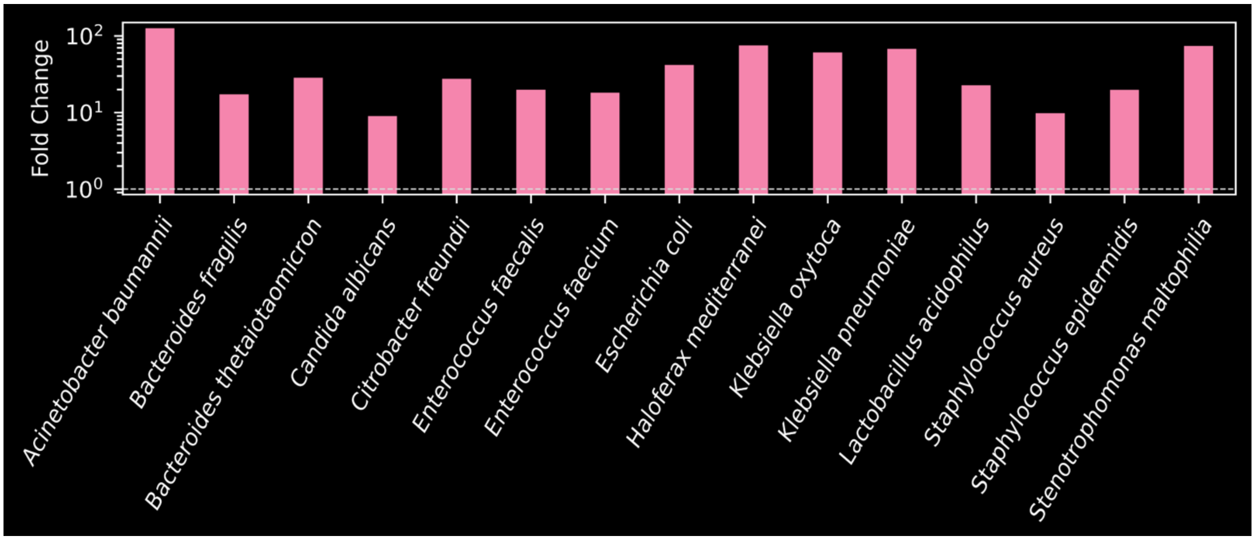

**Figure S4: Color legend of strains used in the consortium for host-microbiome-food interactions.**

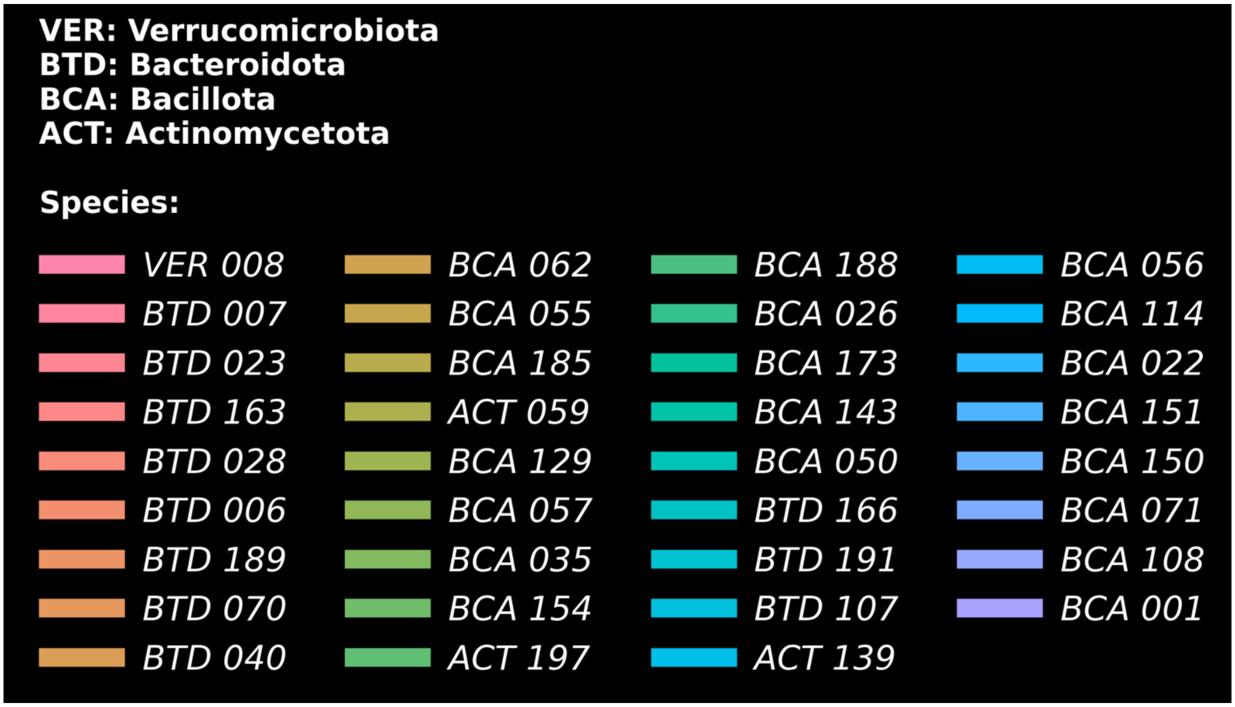


**Figure S5: Abundances measured by HiPR-Map and sequencing in tissue samples.** Comparison of relative abundances measured by HiPR-Map in tissue samples and sequencing of adjacent tissue sections. (Inset) The Spearman rank correlations (ρ) between HiPR-Map and sequencing are calculated using all taxa.

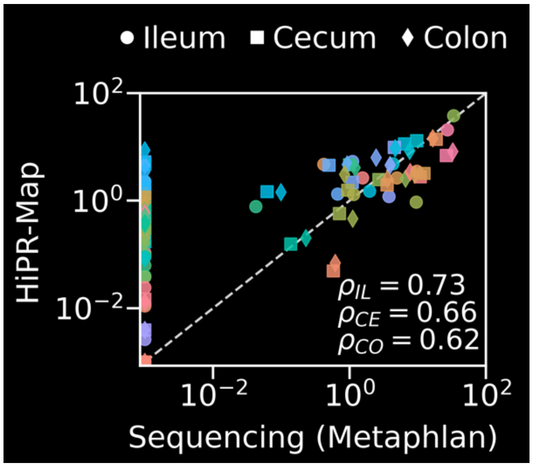

**Figure S6: Distribution of microbes in and around food particles.** Histogram showing the abundance distribution of each taxon relative to the nearest food particle in the ileum, cecum, and colon. The color legend is shown in **Fig. S4**.

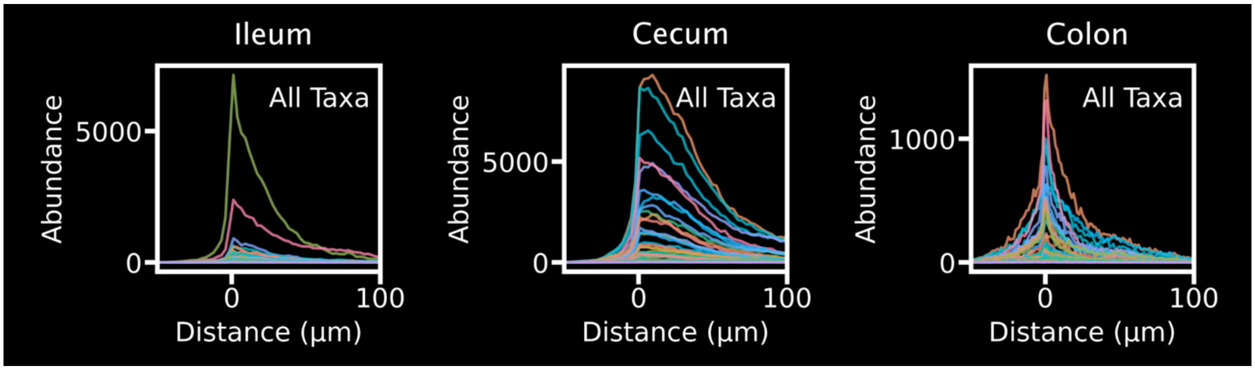


**Figure S7: Reference images of wheat bran.** Wheat bran was prepared and imaged as described in **Methods**. Left: 20X magnification. Right: 63X magnification. Scale bars on both indicate 20 µm.

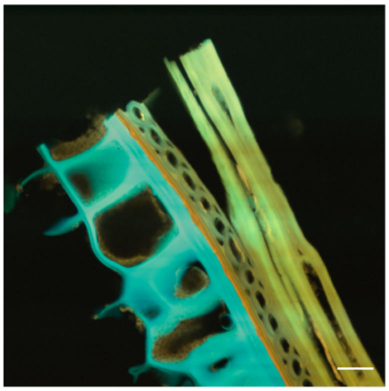

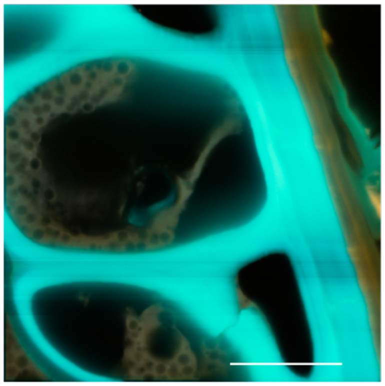
